# Convenient screening for drug resistance mutations from historical febrile malaria samples across Kenya

**DOI:** 10.1101/2025.09.09.675152

**Authors:** John Magudha, Leonard Ndwiga, Mercy Y. Akinyi, Kevin Wamae, Victor Osoti, Regina Kandie, Rosebella Kiplagat, Kibor Keitany, Joel L. Bargul, Hoseah M. Akala, L. Isabella Ochola-Oyier

## Abstract

**Background:** Historically, chloroquine resistance emerged that was driven by mutations in the chloroquine resistance transporter (*Pfcrt)* gene. This led to the global withdrawal of chloroquine in 1998 and its subsequent replacement with sulfadoxine-pyrimethamine, whose efficacy was compromised by a high prevalence of mutations in the dihydrofolate reductase and the dihydropteroate synthase genes by 2004. Consequently, artemisinin-based combination therapies (ACTs) were introduced in 2006. Since then, thirteen mutations in the kelch 13 (Pfk13) propeller domain have emerged and validated by the World Health Organization (WHO) as markers of partial artemisinin resistance. This study aimed to characterize temporal trends in both established, *Pfcrt* and *Pfk13* and less well-described potential markers, cysteine desulfurase (*Pfnfs)* and *Pfcoronin*, using febrile malaria samples collected across diverse regions of Kenya between 2013 and 2022.

**Methods:** The temporal trend of these markers of resistance were assessed by screening archived *P. falciparum* positive dried blood spots (DBS). A total of 1,750 DBS samples collected from Therapeutic Efficacy Studies (TES) conducted in: Kwale (2013, n=350), Kisumu (2015, n=314), Busia (2016, n=334), Kisii (2017, n=314), Kwale (2018, n=150), and a *hrp2* study conducted Kisii (2022, n=288). Parasite genomic DNA was extracted using the Chelex-saponin method and confirmed by a *Pf* 18S RT-PCR. *Pfk13, Pfcrt, Pfnfs* and *Pfcoronin* PCR amplicons were sequenced using capillary electrophoresis, Illumina Miseq or the Oxford Nanopore (GridION) platform.

**Results:** The prevalence of *Pfcrt* mutations declined over time and no WHO validated *Pfk13* mutations associated with artemisinin resistance were detected. However, synonymous substitutions at WHO-validated codons C469C and P553P were identified. In the *PfCoronin* gene, non-synonymous mutations distinct from those reported in West Africa were observed at high frequencies (>75%). Notably, the *Pfnfs*-K65Q mutation, previously associated with reduced lumefantrine sensitivity in West Africa, was detected in over 80% of samples.

Our findings reveal differences in some antimalarial resistance genetic markers between observations made in The Gambia and Senegal (West Africa) and Kilifi (East Africa). Based on the convenient sample set, there were no WHO validated k13 mutations up until 2022, suggesting continued ACT efficacy in Kenya. This study underscores the importance of continued molecular surveillance and suggests that resistance may evolve through different pathways in East compared to West Africa and Southeast Asia.

## Introduction

The persistence of the deadly *Plasmodium falciparum* malaria parasite, despite the roll out of preventive measures and antimalarials for treatment, has led to a global malaria disease burden of 263 million cases in 2023 alone, according to the World Health Organization (WHO) (World Malaria Report, 2024). Efforts to reduce this burden of malaria continue to depend on three main pillars, vaccination, vector control, rapid case detection and the use of effective antimalarials [2].

The WHO recommends the use of a combination of compounds with distinct modes of action for malaria treatment, ensuring effective cure rates to impede the development of resistance. Artemisinin-based combination therapies (ACTs) are the first-line treatment for *P. falciparum* uncomplicated malaria in all endemic countries [3]. ACTs integrate an artemisinin derivative with a non-artemisinin partner drug. Although the efficacy of ACT is dependent on both agents, the artemisinin is critical to reduce the bulk of parasite biomass during the first 3 days of treatment with the partner drug eliminating the remaining parasites [4].

However, areas characterized by low rates of mixed-strain transmission, particularly in Southeast Asia, have been the initial focal points that exhibit resistance to antimalarial drugs [5]. The emergence of resistance to artemisinin and its derivatives in Southeast Asia, manifested as delayed parasite clearance following treatment with these drugs, is of great concern [6]. Recent work attributed the artemisinin resistance phenotype found in Southeast Asia to mutations in PF3D7_1343700, which encodes a protein homologous to kelch proteins from other organisms [7]. Artemisinin-resistant parasites have been shown to harbor multiple mutations in the propeller domain of the Kelch 13 protein. Mutations in *P. falciparum* kelch 13 (*Pfk13*) gene, initially identified in Southeast Asia, including Y493H, R539T, I543T, C580Y [7], have rarely been observed in African countries. In Rwanda, G449A, C469Y and R561H mutations have been reported [8,9], while Uganda has documented C469Y, R561H and A675V mutations [10]. Similarly, Democratic Republic of Congo (DRC) has reported 441I, C469Y and R561J mutations [11]. Sudan, Eritrea and Ethiopia have all reported the R622I mutation [12,13]. Kenya has documented the C469Y, P553L and A675V mutations [14–16], and Tanzania has identified the R561H and R622I mutations [17–19]. These findings suggest a widespread emergence of *Pfk13* mutations which in Ethiopia [20,21], Rwanda [22], Uganda [23,24] and Tanzania [25] have been associated with delayed parasite clearance, raising concerns about the potential of ACT failure across the region [26].

Additional markers linked to chloroquine resistance, the chloroquine resistance transporter (*Pfcrt)*, lumefantrine tolerance potentially cysteine desulfurase (*Pfnfs*) [27] and artemisinin resistance, potentially coronin have been identified [Demas et. al., 2018]. Chloroquine was used as a first-line antimalarial until 1998 when it was retracted as a result of widespread reports of resistance in Sub-Saharan Africa [28]. Chloroquine resistance was primarily attributed to the *Pfcrt* 76T mutation [29]. The cysteine desulfurase (nfs) K65 allele was implicated in lumefantrine resistance in West Africa [30] whereas mutations 50E, 100K and 107V mutations in *Pfcoronin* were associated with artemisinin resistance in Senegal [31].

In this study, the utility of screening therapeutic efficacy studies (TES) samples for known and novel drug resistance mutations was examined. *Pfcrt* was included as it is well known that there has been a reduction in the mutant genotype over time [32,33]. The novel markers, *Pfnfs* and *Pfcoronin*, described as containing potential drug resistance mutations were sampled to define the presence of these mutations previously identified in West Africa. The *Pfk13* mutations were also assessed to provide the historical context from symptomatic samples from Kenya. At the same time, a mix of genotyping tools were used, initially Sanger sequencing, Illumina and due to a shortage of illumina reagents ONT was used. Thus, the utility of using different assays based on reagents availability was done to examine the generation of standard data outputs of SNP frequencies from different platforms.

## Methods

### Study Area and Sampling

Therapeutic efficacy studies (TES) were carried out at four different sites in Kenya, Kwale county in 2013, Kisumu County in 2015, Busia County in 2016 and Kisii county in 2017. A follow-up efficacy trial of AL only was conducted in Kwale county in 2018 [34]. Children aged between 6 months to 14 years seeking treatment at the outpatient departments of participating health facilities with signs of uncomplicated malaria were sampled. Eligible children for whom parental/guardian informed consent was obtained were randomized into a therapeutic efficacy study (TES) to receive dihydroartemisinin-piperaquine (DP) or artemether-lumefantrine (AL). Participants were then required to report to the study clinic on days 1, 2, 3, 7, 14, 21, 28 and 42. Approximately, 0.05ml of blood from a finger prick was collected and used to prepare thin and thick smears as well as to prepare dried blood spots (DBS). Slides were examined and stained with Giemsa under the microscope to detect the presence of malaria parasites and provide parasite density estimates as parasites per microlitre. In 2022, a *hrp2* study was conducted in Kisii County, Kenya, a highland malaria epidemic-prone region. This study was part of a broader cross-sectional study aimed at characterizing Histidine rich protein (HRP) 2/3 gene deletions in patients of all ages presenting with uncomplicated malaria reporting at primary healthcare facilities across 10 high and low malaria transmission counties: Nairobi, Kirinyaga, Homa Bay, Kilifi, Garissa, Kisumu, Kwale, Trans Nzoia and Tana River.

### Parasite genotyping

Parasite DNA was extracted from 1,750 DBS samples using the Chelex-saponin method, according to the manufacturer’s protocol. This comprised of 350 samples from Kwale (2013), 314 from Kisumu (2015), 334 from Busia (2016), 314 from Kisii (2017), 150 from Kwale (2018) and 288 from Kisii (2022). Briefly, samples were lysed with 0.5% (w/v) saponin in 1X Phosphate-buffered saline (PBS) overnight. Following saponin aspiration, the discs were incubated in 1ml 1X PBS at 4°C for 30 minutes after which 150μl of 6% (w/v) Chelex in nuclease free water was added and incubated for 10 minutes at 97°C. The samples were then centrifuged at 4,000 × g for 5 min, and 120μL of the DNA-containing solution was stored at −20°C for sequencing. The parasite DNA was screened for the presence of *Plasmodium falciparum* (*Pf*) using the TaqMan probe-based PCR using *P. falciparum* 18S rRNA forward and reverse primers, as described in (Osoti et al., 2022). Briefly, 2.5µL of each primer (10 pmol/µL), 0.625µL of 18S probe (10 pmol/µL), 12.5µL of 2X TaqMan universal PCR master mix and 6.75µL of sample or 3D7 control samples, with the remaining volume PCR clean water was used. The reaction was done using the following qPCR cycling conditions: 50°C for 2 min, 95°C for 10 min, and then 45 cycles of 95°C for 15 s and 95°C for 1 min. A Ct cut-off of ≥39 was taken as a negative sample.

#### *Pfk13, Pfcrt, Pfnfs* and *coronin* capillary sequencing

Amplicons were generated from genomic DNA for the *Pfk13* gene from the *Pfhrp2* study and the *Pfcrt, Pfnfs*, and coronin genes from the Therapeutic Efficacy Studies conducted in Busia, Kisii, Kisumu and Kwale counties (Table 2:). PCR amplifications were performed using the Expand High Fidelity PCR System (Roche) and HotStarTaq Master Mix (Qiagen, Cat. No. 202602). All reactions were performed to a final volume of 10µl, and each reaction mixture contained 200µM dNTPs, 300nM of each primer, 1X PCR buffer (containing 1.5mM MgCl2, 2.5mM of MgCl2 (buffer 2), 0.035U of Taq and 0.7µl of the template DNA. The cycling conditions of the four gene fragments were as follows: 94°C for 2 minutes, 94°C for 15 seconds, 30 cycles at annealing temperatures shown in Table 1 for each gene(X), 68°C for 2 minutes and a final extension of 7 minutes at 72°C. The amplified PCR products were analyzed on 1.5% (w/v) agarose gel, purified using ExoSAP-IT® (Affymetrix, Santa Clara, CA) as per the manufacturer’s instructions) and then sequenced using the BigDye® Terminator v3.1 Cycle Sequencing Kit reaction mix (Applied Biosystems, USA) using 3730XL sequencer. The raw sequence data from both forward and reverse sequences were assembled into contigs using CLC Main Workbench. This method was utilized for these genes as the loci of interest were known for coronin, *Pfcrt* and *Pfnfs* and for k13 to compare with illumina that can detect low frequency SNPs.

**Table 1:**
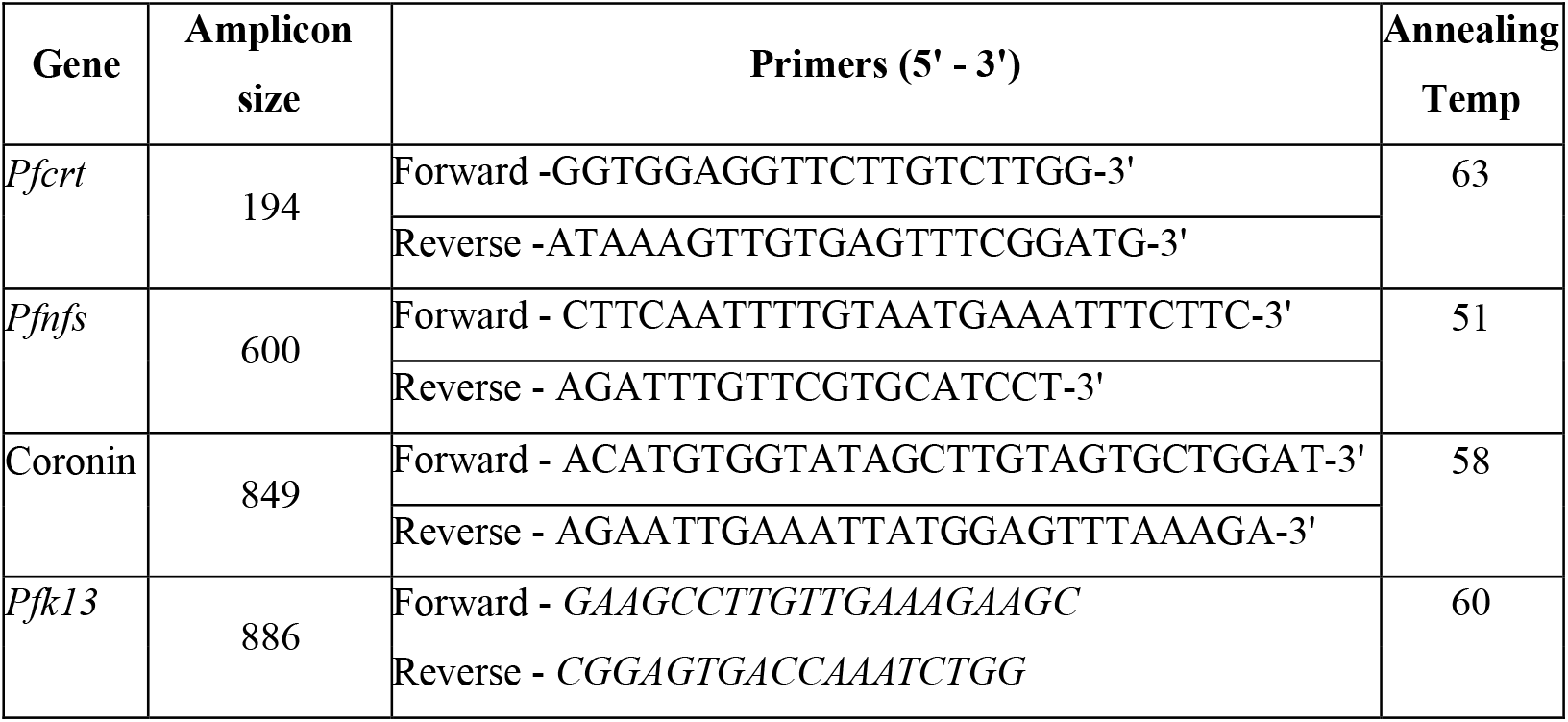
List of Primers used for capillary and amplicon sequencing.

**Table 2:** Sequencing platforms used to genotype *P. falciparum* drug resistance gene amplicons from the different study sites

| Study | Site(s) | Sequencing Platform(s) | Gene(s) Sequenced |
| --- | --- | --- | --- |
| TES | Busia, Kisii, Kisumu, Kwale | Capillary | coronin, nfs, crt |
| TES | Busia, Kisii, Kisumu, Kwale | Illumina | k13 |
| TES | Busia, Kisii, Kisumu, Kwale | ONT | k13 |
| HRP2 | Kisii | Capillary & Illumina | k13 |

#### *Pfk13* drug resistance marker genotyping

Amplicons spanning the codon 425 – 720 of the Kelch 13 (K13: PF3D7_1347700) propeller domain were generated using conventional PCR. The PCR assay was set up as follows: 1µL of template DNA (<25 ng), 0.2µL of Q5 High-Fidelity DNA Polymerase (0.02 U/µL, New England BioLabs), 1µL (10mM) forward primer, 1µL (10mM) reverse primer, 0.4µL (10mM) dNTPs (New England BioLabs), 4µL (5X) Q5 reaction buffer (New England BioLabs), and 12.4µL nuclease-free water. The PCR conditions were as follows: initial denaturation (98°C for 30 seconds), followed by 30 cycles of denaturation (98°C for 10 seconds), annealing (60°C for 30 seconds), extension (72°C for 30 seconds), and final extension (72°C for 2 minutes). PCR products were visualized on 1% (w/v) agarose gels stained with Red Safe Nucleic Acid Staining Solution. PCR products were purified using 1X AMPure Beads (Beckman Coulter, USA), followed by two ethanol (80%) wash steps. Purified PCR products were quantified with Qubit dsDNA HS Assay Kit. Sequencing was performed with either ONT, Illumina or Capillary sequencing as shown in Table 2: below.

For Illumina, Nextera XT (Illumina) and Kapa HyperPrep kits (Roche) were used to add sequencing adaptors and unique barcodes. Transposome was used to fragment and tag DNA with adaptors. A limited-cycle PCR was used to add unique indices to each sample. Using 0.8X AMPure XP beads, libraries of size >300bp were selected and finally, 5.0μl of each library pooled and loaded on the Illumina MiSeq run.

For ONT, End-prep reaction was performed on the purified amplicons using NEBNext Ultra II End repair/dA-tailing Kit (New England BioLabs, USA) and from this, 2μL of DNA was used for barcode ligation using Native Barcoding Expansion 96 (Oxford Nanopore Technology, Oxford, UK) and NEBNext Blunt/TA Ligase Master Mix (New England BioLabs, USA). The barcoded samples were pooled and cleaned using 0.4X AMPure XP beads. Adapter ligation of the cleaned barcoded library was done using 50–100 ng using the NEBNext Quick Ligation Module reagents (New England BioLabs, USA) and Adaptor Mix II (Oxford Nanopore Technology, Oxford, UK). Final clean-up was performed using 125 μL of Short Fragment Buffer (Oxford Nanopore Technology, Oxford, UK). The library was eluted in 15μL Elution Buffer (ONT, Oxford, UK). The final library was normalised to 15–70 ng, loaded on a SpotON R9 flow cell and sequenced on the GridION.

### Sequence Data Analysis

Sequence data from genotyping of the three Sanger-sequenced drug resistance markers was analyzed using CLC Genomics Workbench v23.0. SeekDeep version 3.0.1 was used in analysis of the Illumina [35] and ONT [36] generated data. The paired consensus reads for each sample were trimmed and clustered to estimate the frequency of K13 mutations. The SNP frequencies were calculated using R version 4.2.1. Within-sample frequency was calculated by dividing the number of reads supporting the non-reference allele by the total number of reads per sample.

## Results

### Analysis of Kisii HRP2 survey samples for k13 polymorphisms

A total of 288 samples were screened by 18S rRNA real time-PCR (RT-PCR) with 98/288 (34%) samples being positive with detectable levels of *Pf* DNA. Out of these, 90/98 and 92/98 samples had good-quality *Pfk13* data for Sanger and Illumina sequencing, respectively. Out of the 90 samples analyzed by Sanger sequencing, only one sample was found to harbor both a synonymous mutation (A627A), and a non-synonymous mutation (S679T) within the k13 propeller domain. Illumina sequencing only identified the A627A and not the non-synonymous mutation (S679T). Specifically, a total of 5 synonymous mutations were identified in 15/92 samples (16.3%) sequenced by Illumina platform (Supplementary Tables

Table). The prevalence of the P553P mutation was highest in 8/92 samples followed by the C469C in 3/92 samples and A427A in 2/92 samples. The remaining synonymous mutations (G548G and A627A) were observed in 1 sample each. The G548G and A627A synonymous mutations had a >98% within sample frequency (Table S1). Illumina identified more low frequency k13 alleles than capillary sequencing.

#### Prevalence of *k13* mutations in the TES samples

A total of 6 mutations were identified in the K13 gene, with only the A578S being non-synonymous. The A578S mutation was identified in both Kisii and Busia at a frequency of 0.3% [1/288] and 1.4% [1/72], respectively (Table 4). Two WHO validated artemisinin resistance associated mutations were observed though the mutations were synonymous, C469C in Busia 2016 and Kisii in both 2017 and 2022 and P553P in Kisii 2022. All the mutations were low frequency (<5%) except the P553P synonymous mutation.

**Table 4:** The frequency of mutations in the *Pfk13* gene

| Year | County | Genotyping Assay | Samples Genotyped | Allele | Mutant Allele Frequency, n [%] |
| --- | --- | --- | --- | --- | --- |
| 2013 | Kwale | ONT | 57 | None | 0 |
| 2015 | Kisumu | ONT | 140 | A578S | 3 [2.1] |
| 2016 | Busia | Illumina | 72 | **C469C** | 1 [1.4] |
|  |  |  |  | A578S | 1 [1.4] |
|  |  | ONT | 43 | None | 0 |
| 2017 | Kisii | Illumina | 286 | **C469C** | 2 [0.7] |
|  |  |  |  | R471R | 1 [0.3] |
|  |  |  |  | A578S | 1 [0.3] |
|  |  |  |  | G638G | 1 [0.3] |
|  |  |  |  | V666V | 3 [1] |
|  |  |  |  | N694N | 2 [0.7] |
| 2018 | Kwale | ONT | 37 | None | 0 |
| 2022 | Kisii | Illumina | 92 | A427A | 2 [2.17] |
|  |  |  |  | **C469C** | 3 [3.3] |
|  |  |  |  | G548G | 1 [1.1] |
|  |  |  |  | **P553P** | 8 [8.7] |
|  |  |  |  | A627A | 1 [1.1] |
| 2022 | Kisii | Capillary | 90 | A627A | 1 [1.1] |
|  |  |  |  | S679T | 1 [1.1] |

n is the number and [ percentage] of isolates carrying each mutation. None and 0 indicate no mutations were identified.

**Table 5:** The frequency of mutations in CRT, NFS and Coronin by Capillary Electrophoresis

| Gene | crt %**[n]** |  |  |  | coronin %**[n]** | nfs %**[n]** |  |  |  |  |  |  |
| --- | --- | --- | --- | --- | --- | --- | --- | --- | --- | --- | --- | --- |
| Nucleotide | *G222T* | *T225A* | *A227C* | *T248A* | *A547G* | *GI85A* | *A193C* | *A200G* | *A304G* | *C328G* | *G359T* | *G390C* |
| Codon | *M74I* | *N75E* | *K76T* | - | *S183G* | *S62N* | *K65Q* | *E67G* | *N102D* | *Q110E* | *S120I* | *E130D* |
| Kisumu 2015 | 1.7 [3] | 1.7 [3] | 1.7 [3] | - | 88.5 [46] | 88 [66] | 82.7 [62] | 84 [63] | 1.3 [1] | 14.7 [11] | 42.7 [32] | 20 [15] |
| Busia 2016 | 2.7 [4] | 3.3 [5] | 4.7 [7] | - | 75 [24] | 85.4 [41] | 89.6 [43] | 89.6 [43] | - | - | 20.5 [8] | 6.1 [2] |
| Kisii 2017 | 0 [0] | 0 [0] | 0 [0] | 1.2 [1] | 69.7 [23] | 86.9 [119] | 82.7 [115] | 83.5 [116] | 0.7 [1] | 12.2 [18] | 38.9 [58] | 24.5 [37] |
| Kwale 2018 | 0 [0] | 0 [0] | 0 [0] | - | 80 [32] | 85.4 [76] | 86.9 [73] | 74.4 [67] | 0 [0] | 8.9 [8] | 35.6 [32] | 27.8 [25] |

The total number of samples sequenced for crt (Kisumu, n=179; Busia, n=153; Kisii, n=84; Kwale, n=89), for coronin (Kisumu, n=52; Busia, n=52; Kisii, n=34; Kwale, n=87), and for nfs (Kisumu, n=74; Busia, n=51; Kisii, n=160; Kwale, n=89).

#### Prevalence of other drug resistance mutations in the TES samples

A total of 3 genes, *Pfcrt, Pfnfs* and coronin were genotyped to determine the frequency of known single nucleotide polymorphisms and *coronin* to identify any novel mutations across the four sites. The K76T mutations identified in *Pfcrt* gene was highest in Busia 2016 at a frequency of 4.7% followed by Nyando 2015 at 1.7%. There were no *Pfcrt* mutations identified in both Kisii (2017) and Kwale (2018). A number of mutations were identified in the *Pfnfs* gene including the 65Q allele. Only one mutation 183G, in the *coronin* gene was identified in all the four sites, **Table 5**.

### Comparison of the ONT and Illumina generated reads

Following Pfk13 gene amplifications, a total of 354 samples were successfully sequenced by Illumina, including 68 from Busia and 286 from Kisii. Additionally, 277 samples were successfully sequenced by ONT, comprising a different set of 43 samples from Busia, 94 from Kwale (2013 and 2018), and 140 from Kisumu. Illumina combined all the samples sequenced into one sequencing run generating a total of 42 million reads, whereas the samples sequenced on ONT were distributed into three runs resulting in 21 million, 14 million and 15 million reads. Though it is not a head-to-head comparison of platforms as no samples were run on both platforms, overall, the number of reads per sample (Figure 1A) was highest from Illumina. Kisii had the highest number of K13 reads generated where the largest number of samples were primarily sequenced by Illumina.

**Figure 1.**
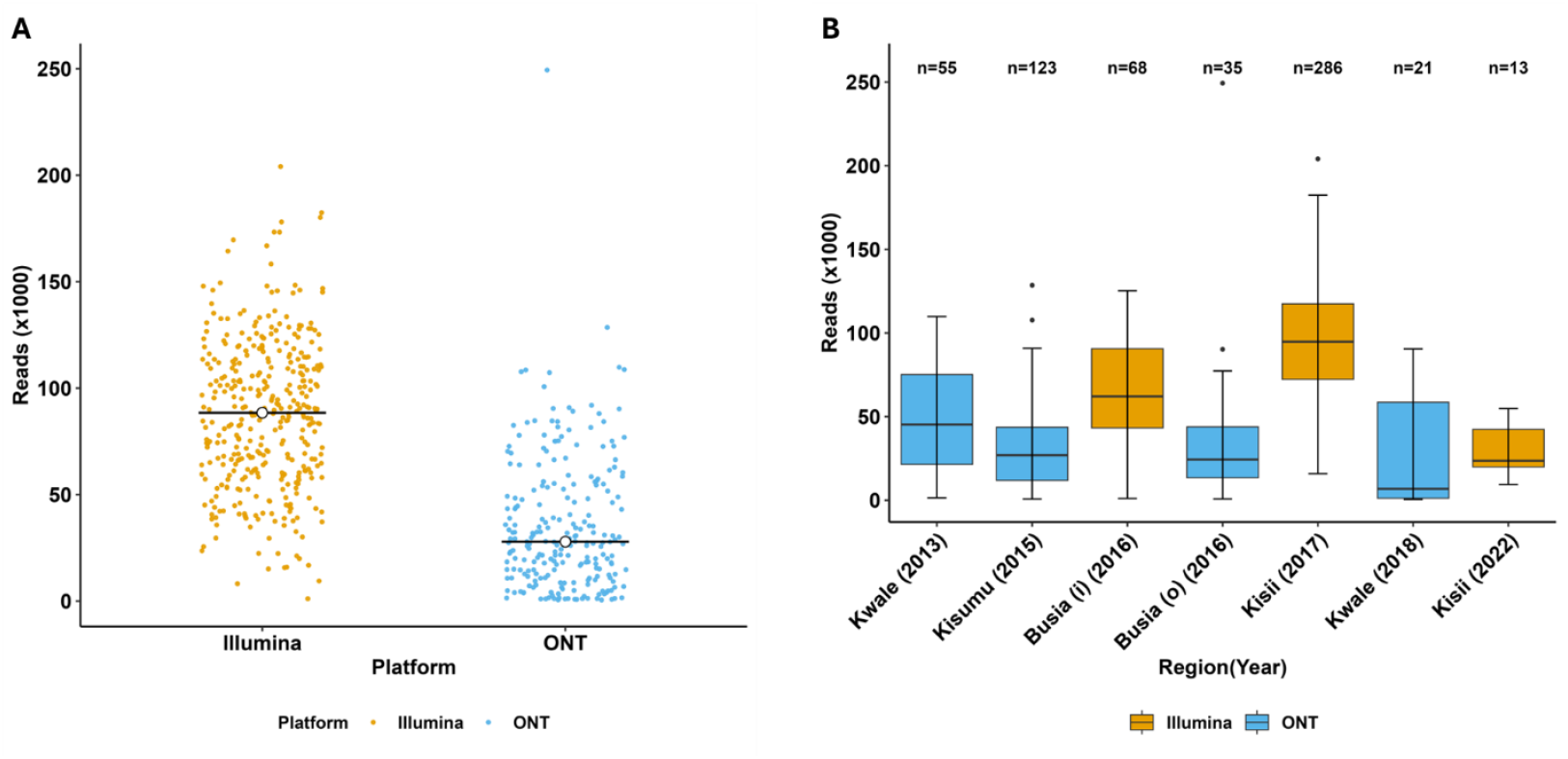
Total number of reads generated by platform. **A)** The total number of k13 reads generated from the Illumina-MiSeq and ONT sequencing platforms: The mean read count was 87,686 and 35,906 for Illumina and ONT, respectively. **B)** k13 reads distribution across sites and platforms: The median read counts were: 45248 (Kwale 2013 - ONT), 6835 (Kwale 2018 - ONT), 26925 (Kisumu 2015 - ONT), 62068 (Busia 2016 - Illumina), 24355 (Busia 2016 - ONT), 94791 (Kisii 2017 - Illumina) and 23612 (Kisii 2022 - Illumina).

## Discussion

The propensity for the *Pfk13* gene to accumulate mutations potentially appears initially as synonymous mutations generating the basic working material for continued natural selection. This early data provides evidence of important mutational changes identified by molecular surveillance. These changes provide a warning of a changing gene over time, accumulating possible detrimental mutations that can result in resistance conferring variants. Additionally, the mutations are observed at low frequency in a few samples in mixed genotype infections. Although synonymous mutations have traditionally been regarded as functionally neutral, emerging evidence suggests they may play a more active role in shaping evolutionary trajectories. In yeast, Shen et al., (2022) demonstrated that synonymous mutations can significantly reduce fitness by disrupting mRNA levels, thus affecting gene expression [37]. Similarly, Faheem et al. (2021) showed that such mutations can alter mRNA 3D structure, influencing stability and translation efficiency. These changes may set the stage for adaptive non-synonymous mutations to emerge. Forman and Coller (2022), proposed that synonymous mutations may facilitate evolutionary access to previously unfavorable amino acid changes by modifying codon usage. In our study, we observed synonymous mutations at k13 positions C469C and P553P, sites where non-synonymous variants associated with artemisinin resistance (e.g., C469Y and P553L) are now increasingly being reported in the East Africa [38–42]. Recent studies from Kenya have reported non-synonymous mutations emerging at these same loci, such as C469Y in symptomatic and asymptomatic populations and P553L in asymptomatic populations, that are associated with artemisinin resistance [43,44]. This pattern raises the possibility that drug-resistant *P. falciparum* parasites could emerge through random synonymous substitutions, which subsequently evolve into non-synonymous mutations that are advantageous under continued drug pressure. Previous data has shown that as the mutations rise in high frequency due to positive selection, mutant only infections are observed, as is the case for dhfr (codons 108 and 51) and dhps (codons 437 and 540) mutations [45].

This underscores the need for molecular and phenotypic characterization of the role of synonymous mutations in resistance development. A recent Cameroonian study employed *in silico* modelling and physicochemical property predictions to evaluate the impact of novel k13 mutations and found that at least one variant, L663P, significantly altered the putative stability and conformation of the *Pfk13* propeller domain [46]. This finding underscores the utility of computational approaches in predicting functional and structural impact of novel *Pfk13* mutations implicated in resistance development.

Interestingly, no k13 mutations were observed in the Eastern part of the country, which could be attributed to the low sample size and the potential lower sensitivity of the ONT platform to detect low frequency variants, based on the lower read coverage. Nevertheless, k13 mutations are emerging in Western Kenya, further corroborated by previous publications [44,47,48], highlighting this area as the region to watch and intensify surveillance efforts for k13 mutations and potential artemisinin treatment failure. The recent publications of the presence of validated k13 mutations raise the urgency of a TES evaluation, even though the current evidence is that ACTs remain effective in the country.

The crt mutations provide further evidence of the shift in directional selection to a fully wild-type genotype, with the reduction to zero over time in Kisii and Kwale after 2016. This is in line with previous data from Kisumu [49] and Kilifi [33]. In contrast, the mutations observed in *Pfnfs* and *Pfcoronin* maintain a high frequency across all sites over time. Thus, the Kenyan parasite population sampled is in contrast to West Africa, where there was an observed increase in frequency over time for *Pfnfs* [27] and a completely different mutation from what has been published previously for coronin, codon 183 [50,51]. Though there is evidence to suggest these 2 genes may be involved in lumefantrine and artemisinin resistance, respectively, in some countries in West Africa, they do not appear to be relevant markers in East Africa. Furthermore, the Kenya parasite population are now predominantly wildtype at the Pfcrt locus (CVMNK) compared to parasites from West Africa where there is still a high prevalence of the resistant haplotype CVIET [52,53]. This is not surprising since the drug pressure in East and West Africa is different and genetically the parasite populations are different [27].

This study was not designed as a direct head-to-head comparison of sequencing platforms; however, our results reveal a statistically significant difference in read depths between Illumina Miseq and Oxford Nanopore Technology (ONT) platforms (p < 2e−16). As expected, Illumina consistently produced higher k13 read depths, suggesting enhanced high-throughput sequencing capacity and base-calling accuracy [54]. This improves the ability to detect low frequency changes, however greater accuracy and quality control steps are required to determine a true low frequency variant from an error. In contrast, ONT yielded markedly lower read counts, with considerable variability across years and regions. Nonetheless, the adoption of ONT remains promising due to its cost-effectiveness, portability, and real-time data generation. Further optimization and validation of ONT-based workflows will be crucial to support its integration into national malaria molecular surveillance programs. Sanger sequencing was easier to conduct and analyze, however it primarily detects the dominant genotypes and high frequency mixed genotypes, rare variants were difficult to define. Though the platforms are different, the SNP frequency data generated allowed for comparisons of the same genetic loci across samples from different regions, with the caveat that a lower sample size impacts read coverage.

We acknowledge the limitations inherent in utilizing convenient archived samples. Archived samples are subject to variable storage conditions and durations between collection and processing, which may compromise DNA integrity and yield and influence the variability in sequencing outcomes. The use of convenient samples from different geographies and time points could bias the interpretation of population-level patterns. Practical considerations with regards to platform reagent availability and costs influenced the choice of samples for sequencing. Consequently, certain sites and time periods were disproportionately represented across sequencing platforms, limiting direct comparability.

Despite the above limitations, our approach provided a broader temporal and geographical perspective of *P. falciparum k13* polymorphisms across Western and Coastal Kenya between 2013 and 2022, which would not have been possible through a single study site or assay. To mitigate these challenges, data was harmonized across studies, consistent analytical pipelines utilized and highlighted methodological or sampling differences that could influence interpretation. We also included Pfcrt, a known antimalarial drug resistance marker with extensive historical data to support the interpretation of the observed trends.

This study offers valuable insights into the evolving genetic landscape of *P. falciparum* k13 antimalarial resistance in Western and Coastal Kenya between 2013 and 2022. The gradual increase of the C469C mutation in Kisii may represent synonymous mutations that precede the emergence of resistance-conferring variants. In addition, it highlights the geographical difference between East and West Africa with the high prevalence of *Pfnfs* and no *Pfcoronin* mutations, suggesting different mechanisms for the emergence of resistance between the different regions of Africa. This retrospective look demonstrates that continuous molecular surveillance remains crucial for the early detection of emerging resistance markers.

## Authors’ contributions

LIOO secured funding, conceived the study, and designed the experiments. JM and LN performed the experiments. JM, LN, MA, KW, and LIOO analyzed the data. LIOO, RK, RK, KK, and HA supervised fieldwork and were responsible for sample acquisition. JM, LN, and LIOO drafted the initial manuscript. All authors (JM, LN, MA, KW, VO, RK, KK, JL, HA, and LIOO) contributed to data interpretation, critically reviewed the manuscript, and approved the final version

## Acknowledgements

We are grateful to the study participants and their guardians, as well as the field and laboratory teams involved in this work. We also thank the National Malaria Control Programme (NMCP) for coordinating the Therapeutic Efficacy Studies (TES). All laboratory experiments were carried out at the KEMRI–Wellcome Trust Research Programme laboratories.

## Supplementary Tables

**Table S1.** Within sample non-reference (mutant) k13 allele frequencies in samples collected from the Kisii hrp2/3 study.

| codon position | nucleotide position | reference allele | non-reference allele | Total number of reads, within sample<br>non-ref allele frequency (%) |
| --- | --- | --- | --- | --- |
| A427A | 1281 | C | A | 115 (6) |
|  | 1281 | C | A | 115 (6) |
| C469C | 1407 | C | T | 22374 (27) |
|  |  | C | T | 22322 (26) |
|  |  | C | T | 23612 (27) |
| P553P | 1659 | G | A | 45083 (6) |
|  |  | G | A | 15101 (6) |
|  |  | G | A | 548181 (5) |
|  |  | G | A | 30103 (5) |
|  |  | G | A | 40158 (7) |
|  |  | G | A | 19947 (6) |
|  |  | G | A | 53047 (6) |
|  |  | G | A | 9440 (11) |
| G548G | 1644 | C | T | 15702 (98) |
| A627A | 1881 | T | A | 42347 (99) |

This table shows the synonymous mutations identified in 15 *P. falciparum* isolates using illumina AmpSeq. For each mutation, the total number of sequencing reads supporting the non-reference allele is shown alongside its proportion (%) of the total reads at that position. Each row represents a different sample.

